# In vivo deciduous dental eruption in LuiKotale bonobos and Gombe chimpanzees

**DOI:** 10.1101/2021.01.06.425588

**Authors:** Sean M. Lee, L. J. Sutherland, Barbara Fruth, Carson M. Murray, Elizabeth V. Lonsdorf, Keely Arbenz-Smith, Rafael Augusto, Sean Brogan, Stephanie L. Canington, Kevin C. Lee, Kate McGrath, Shannon C. McFarlin, Gottfried Hohmann

## Abstract

Existing data on bonobo and chimpanzee dental eruption timing are derived predominantly from captive individuals or deceased wild individuals. However, recent advances in noninvasive photographic monitoring of living, wild apes have greatly expanded our knowledge of chimpanzee dental eruption in relatively healthy individuals under naturalistic conditions. We employ similar methods to expand on this knowledge by reporting deciduous dental eruption ages in living, wild bonobos and chimpanzees from LuiKotale, Democratic Republic of the Congo and Gombe National Park, Tanzania, respectively. Deciduous dental eruption ages in our sample generally fall within the range of variation previously documented for captive chimpanzees. We also found substantial variation in deciduous canine eruption timing, particularly among bonobos. One bonobo had a deciduous canine present by 227 days old while another did not have a deciduous canine present at 477 days old. As more data accumulate from these populations, future studies should consider sources of variation in deciduous canine eruption timing and relationships with other aspects of life history.

## Introduction

Dental eruption is typically defined as the process of tooth movement from within the alveolar crypt to the fully occluded position in the dental arcade (Kuykendall, Mahoney, and Conroy 1992; Nissen and Riesen 1964). Early comparative studies highlighted covariance between dental eruption timing and life history milestones across a wide range of primates, most notably documenting an association between the timing of first permanent molar eruption and weaning (Godfrey et al. 2001; Smith 1989, 1992). Thus, age at permanent tooth eruption, which can be calculated from incremental growth features in teeth of immature individuals, has been an important proxy for inferring hominin life history (e.g., Kelley and Schwartz 2012; Bromage and Dean 1985). However, other studies questioned the utility of extant great ape dental eruption timing for the purpose of inferring hominin life history (Kelley, Schwartz, and Smith 2020; Machanda et al. 2015; Smith et al. 2013; reviewed in Robson and Wood 2008). Thus, there is a need for more data within narrower phylogenetic contexts, e.g., within species and genera, to generate a better understanding of the relationship between dental eruption and life history milestones (see Kelley et al., 2020).

Until recently, information on dental eruption in our closest living great ape relatives, bonobos (*Pan paniscus*) and chimpanzees (*P. troglodytes*), was based predominantly on living or deceased captive individuals (Bolter and Zihlman 2011; Conroy and Mahoney 1991; Kraemer et al. 1982; Kuykendall, Mahoney, and Conroy 1992; Nissen and Riesen 1964; Smith, Crummett, and Brandt 1994) or deceased free-ranging individuals of chronological age that was either known from living observations or determined from dental histology (Smith and Boesch 2011; Zihlman, Bolter, and Boesch 2004, 2007; Kelley, Schwartz, and Smith 2020). Pusey (1978) described dental eruption observations in a small number of living, free-ranging chimpanzees from the Gombe East African chimpanzee population, and more recently, Smith et al. (2013) and Machanda et al. (2015) have pioneered methods to systematically document dental eruption timing in living, free-ranging chimpanzees. The method developed by Smith et al. (2013) and Machanda et al. (2015) entails photographing the open mouths of individually recognized animals with known life histories to evaluate their stage of dental eruption. This method has substantially improved our knowledge of dental eruption timing and its relationship to other life history variables in free-ranging chimpanzees. However, the studies by Smith et al. (2013) and Machanda et al. (2015) are based on one community of East African chimpanzees from the Kanyawara population. Additional data based on multiple populations and closely related species are needed to evaluate patterns of dental development in hominoids.

Here we report the presence/absence of deciduous teeth in living, wild bonobos from LuiKotale, Democratic Republic of the Congo and East African chimpanzees (*P. t. schweinfurthii*) from Gombe National Park, Tanzania. While most research has focused on permanent dental eruption timing, deciduous tooth eruption may be an important correlate of early infant development that reflects species differences in life history strategies and/or interindividual variation in developmental rates (Mahoney 2019), given potential relationships between deciduous dental eruption and factors such as diet, body size, and/or phylogeny (Smith et al. 2015). We employed a photographic and video scoring method similar to that of Smith et al. (2013) and Machanda et al. (2015). Our results expand on the existing body of data on in vivo dental eruption in chimpanzees and provide the first such data in wild bonobos. These data thus provide critical benchmarks to compare to future and existing data on *Pan* dental eruption and further evaluate the utility of extant great ape dental eruption timing in hominin life history reconstruction.

## Methods

### Study populations

We collected data on bonobos from the Bompusa East and Bompusa West communities at LuiKotale, Democratic Republic of the Congo. We collected data on chimpanzees from the Kasekela community at Gombe National Park, Tanzania. All bonobos and chimpanzees in our study were habituated to human observation. We focused our study on individuals younger than two years of age, as previous studies indicate that in general all deciduous teeth erupt during this period of infancy (e.g., Kuykendall et al. 1992). We determined the birthdate of individuals by taking the midpoint date between the date when the individual’s mother was last observed without the newborn individual (hereon “earliest possible birthdate”) and the date when the mother was first observed with the newborn individual (hereon “latest possible birthdate”). If the earliest and latest possible birthdates were consecutive dates, we assigned the birthdate as the latest possible birthdate. If the number of days between the earliest and latest possible birthdates was an even number of days, we used the later of the two midpoint dates. For all but one individual in our sample, the earliest and latest possible birthdates were separated by less than 24 days. For one bonobo, the earliest and latest possible birthdates were separated by 46 days. Therefore, the largest possible error associated with any individual’s birthdate is 23 days. Our sample includes nine female bonobos, six male bonobos, three female chimpanzees, and four male chimpanzees.

### Photograph and video data collection

At LuiKotale, researchers following bonobos for behavioral data collection collected dental photographs of target individuals opportunistically between April 2014 and July 2017. At Gombe, researchers following chimpanzees for behavioral data collection began collecting dental photographs and video footage beginning in June 2013; this data collection is ongoing and data for this study include photographs and video footage until December 2019. At both study sites, photographs were collected using a high-resolution digital single-lens reflex camera and large aperture telephoto zoom lens. For dental photographs, we scored the presence/absence of teeth directly from the photographs (see next section). For video footage, we used Adobe Premiere to extract photographic stills from video footage when target individuals’ teeth were scorable, i.e., when one can clearly determine which teeth are present, then scored the presence/absence of teeth from these video stills as we describe below for photographs.

### Photograph and video still dental scoring

S.M.L and S.C.M. scored the presence or absence of all teeth from photograph and video still sessions; here, we define a session as one photograph or video still of a given individual or multiple photographs or video stills of a given individual taken on the same day. We used the following coding scheme to score teeth: Present = any part of the tooth crown is visible above the gumline (i.e., from the point of emergence of the cusp tip through the gingival margin, to full functional occlusion); Absent = the tooth is not visible (i.e., the cusp tips have not broken through the gumline); or Z = could not determine because the view of the tooth is obstructed (see Figure 1 for examples). We did not differentiate between the stage of eruption because we felt that Present/Absent was a less subjective metric given that some photographs and stills in our sample were difficult to score accurately beyond Present/Absent (e.g., very pixelated and/or dark). We viewed photographs in Adobe Photoshop and we adjusted image parameters (e.g., brightness) as needed to better assess tooth types and scores. S.M.L. and S.C.M. scored the full dataset by each scoring one subsample of the full dataset. S.M.L. and S.C.M. scored 41 of the same sessions to assess interobserver reliability. We calculated Cohen’s Kappa in R version 4.0.2 (R Core Team 2020) and RStudio version 1.3.1 (RStudio Team 2020) to assess interobserver reliability using the kappa2 function in the irr package (Gamer et al. 2012) and found that Kappa = 0.995. S.M.L. and S.C.M. disagreed on one tooth from one session: where S.M.L. scored Present, S.C.M. scored Z. We reassessed this session and decided to score the tooth Z. After excluding sessions for which we scored all teeth as Z, our sample included 76 bonobo sessions and 16 chimpanzee sessions.

**Figure 1.**
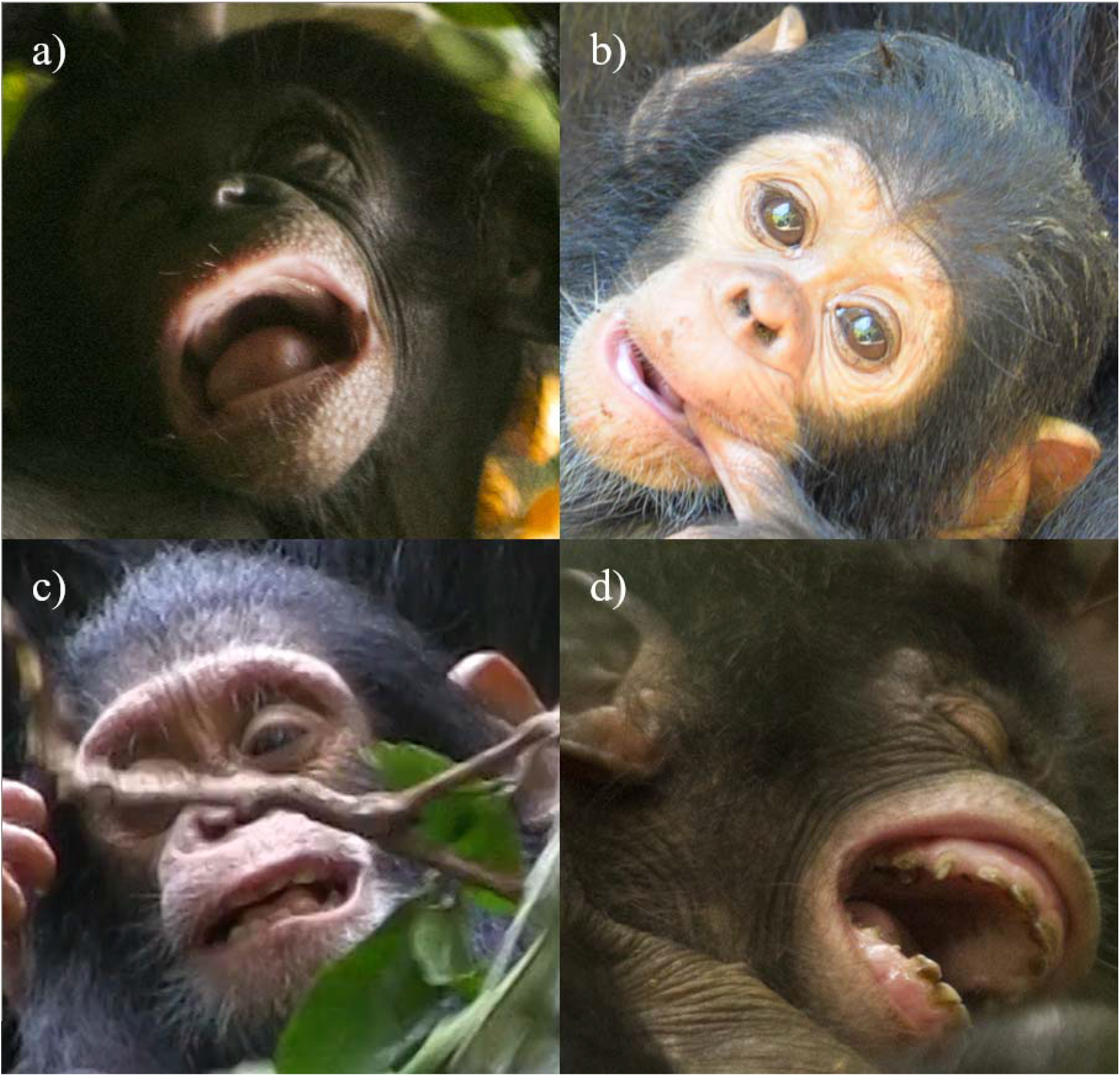
Examples of photographs and video stills. a) Photograph of bonobo (individual PIP at 85 days old) with all maxillary teeth, mandibular incisors, and mandibular canines scored as Absent, and mandibular premolars scored as Z (i.e., could not determine); b) photograph of chimpanzee (GDL at 123 days old) with mandibular first incisors scored as Present, mandibular second incisors and canines scored as Absent, and all other teeth scored as Z; c) video still of chimpanzee (GOS at 371 days old) with mandibular incisors, maxillary incisors, and left mandibular canine scored as Present and all other teeth scored as Z; d) photograph of bonobo (WAT at 227 days old) with several teeth scored as Present, most notably maxillary and mandibular right canines.

We aggregated raw scores in two ways: first, we used a single score for left and right sides of the mouth; thus, if we scored Present or Absent for a given tooth on one side and Z on the other side, we used the Present or Absent score, respectively; if we scored Present on one side and Absent on the other side, we used the most dentally advanced score (i.e., Present). Second, we used a single score for each session; thus, if we scored Present or Absent for a given tooth in one photograph or video still and Z for another photograph or video still from the same individual on the same day, we used the Present or Absent score, respectively.

In our results, we included data for mandibular deciduous teeth from the Kanyawara population of East African chimpanzees, which we estimated from Machanda et al. (2015; fig. 2); these Kanyawara datapoints represent the age of the youngest individual for which emergence of the tooth was observed or the age of the youngest individual for which the tooth was observed to be past emergence in cases when emergence was not directly observed. We did not include maxillary data from the Kanyawara chimpanzee population because Machanda et al. (2015) did not report deciduous eruption ages for maxillary teeth. We also included captive chimpanzee deciduous emergence age ranges from Kuykendall et al. (1992).

**Figure 2.**
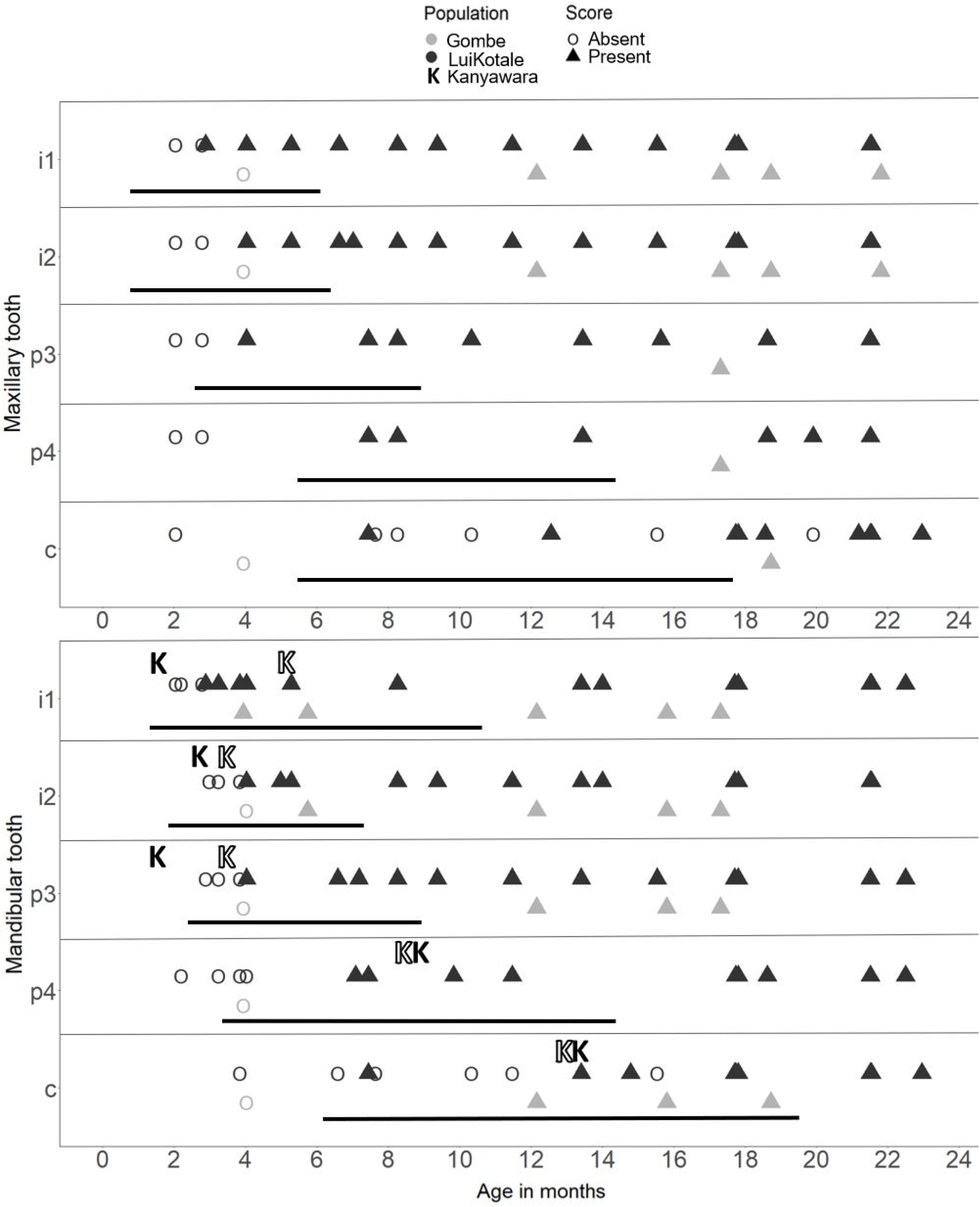
Presence/absence of maxillary and mandibular deciduous teeth. For each tooth representing Gombe and LuiKotale data from the current study, an open circle represents a session for which we scored the tooth as Absent, and a filled triangle represents a session for which we scored the tooth as Present. We utilized criteria to avoid repeated and irrelevant observations for a given individual and tooth: where we have multiple longitudinal sessions of an individual, we only included the last session for which we scored a given tooth as Absent and the first session for which we scored that tooth as Present. Importantly, due to uneven sampling of individuals, filled triangles do not necessarily indicate that the tooth recently erupted in that individual, as they could also indicate that the individual was not sampled regularly prior to the first time that we scored the tooth as Present in that individual. i1 = incisor 1; i2 = incisor 2; c = canine; p3 = premolar 3; p4 = premolar 4. Black horizontal bars represent age ranges for tooth emergence in 41-51 captive chimpanzees from Kuykendall et al. (1992; table 2). Data from Kanyawara (denoted with ‘K’ above) are from Machanda et al. (2015; fig. 2): filled Ks represent age at emergence or youngest age when the tooth was observed to be past emergence in cases when emergence was not directly observed and open Ks indicate the oldest age for which the tooth was observed to be Absent.

The ideal standard for studies of dental eruption timing would be longitudinal observations of tooth emergence from a large sample of subjects conducted systematically at short intervals (i.e., on a daily basis), and over the full duration of development. Or, failing this, sufficient sampling to allow for the calculation of age at emergence for each population using, for example, cumulative distribution functions (see Smith 1991). However, the current study does not meet these criteria. The sample is small and opportunistic, observations were conducted at irregular and often extended intervals, and some ages are poorly represented for any given tooth type. Nevertheless, these data are still extremely valuable given the dearth of data on deciduous dental eruption in wild apes.

### Ethical approval

All methods used in the field were noninvasive and were approved by the Institut Congolaise pour la Conservation de la Nature and the Tanzania Wildlife Research Institute. All aspects of the study comply with the ethics policy of The George Washington University Office of Animal Research Policies and Procedures (https://animalresearch.gwu.edu/) and the guidelines for the ethical treatment of nonhuman primates of the Max Planck Institute for Evolutionary Anthropology (https://www.eva.mpg.de/primat/ethical-guidelines.html).

## Results

The youngest bonobo and chimpanzee that we scored was 62 days old and 120 days old, respectively. Figure 2 illustrates aggregated Present/Absent scores by tooth type and population for maxillary and mandibular teeth. Supplementary Table 1 includes age ranges for the oldest age at which a tooth was scored as Absent and the youngest age at which it was scored as Present for each population. Supplementary Table 2 and Supplementary Table 3 include aggregated raw scores for LuiKotale and Gombe, respectively.

## Discussion

This is the first study to our knowledge that reports dental eruption timing in living, wild bonobos. As such, this study expands our understanding of great ape life history and provides a benchmark with which to compare to future dental eruption studies on wild great apes. Furthermore, this study contributes to the limited body of dental eruption data from living, wild chimpanzees.

Our data for both bonobos and chimpanzees are consistent with the range of variation found in captive chimpanzee deciduous dental eruption timing (Kuykendall et al., 1992). This suggests that the two species exhibit largely similar patterns of deciduous dental eruption timing. However, the range of variation is quite broad; for example, even the incisors, the least variable tooth type, varied in eruption timing by a minimum of approximately four months in the captive chimpanzee study by Kuykendall et al. (1992). Given that all deciduous teeth in this study erupt before the end of two years of age in both species, this variation is considerable and appears to represent a general pattern of high variation in deciduous dental eruption (e.g., see Aiello, Montgomery, and Dean 1991).

The most striking result from our study is the high variation in deciduous canine eruption timing. The youngest bonobo in our sample with a canine present was WAT at 227 days old, while the oldest bonobo yet to have a canine was PIP at 477 days old. The youngest chimpanzee with a canine present was GOS at 371 days old, but we again acknowledge the large gaps in our sampling interval, particularly during the early ages of life. The earliest age of mandibular canine emergence in the Kanyawara chimpanzee sample was approximately 440 days (Machanda et al., 2015), and approximately 186 days in the captive chimpanzee study by Kuykendall et al. (1992). This large variation in deciduous canine emergence ages suggests that deciduous canines may be an important correlate of within and/or between species variation in infant development. Importantly, Smith et al. (1994) show that the range of deciduous canine emergence ages for humans are much later than the range of emergence ages for chimpanzees. Future studies should explore relationships between variation in deciduous canine emergence ages and ecological parameters, maternal characteristics, and other aspects of infant development.

It is important to note that the bonobo in our sample with the canine present at 227 days old, WAT, had earliest and latest possible birthdates separated by 46 days. As we described in the methods, we calculated the birthdate as the midpoint of this range; thus, the error associated with WAT’s birthdate is 23 days. Using the earliest and latest possible birthdates for WAT, his canine would have been present at 204 or 250 days of age, respectively. It is also important to note that we did not consider whether a tooth may have erupted but then was subsequently lost. While this may be unlikely, our criteria for Absent assumes that the tooth has not yet emerged.

Our data indicate broad similarities between species and between captive versus wild conditions. However, our data also underscore the high degree of potentially salient interindividual variation in eruption timing found in previous studies. While we could not investigate sources of such variation given sample size constraints, high interindividual variation in deciduous canine eruption timing in particular represents an important subject of continued study as data from more populations become available. Specifically, more systematic observations at shorter observation intervals and over a larger sample of individuals throughout development are needed to generate statistical treatments regarding the relationship between interindividual variation in emergence ages and other variables, such as sex and maternal characteristics. Additionally, in vivo data from West and Central African chimpanzee populations, as well as other bonobo populations, are needed.

## Author contributions

S.M.L. and G.H. conceived the study; S.M.L., K.A-S., R.A., S.B., S.L.C., K.C.L., E.V.L., K.M., S.C.M., C.M.M., and L.J.S. curated data; S.M.L. wrote the first draft of the manuscript; all authors contributed to subsequent editing, with notable contribution and discussion from B.F., G.H., E.V.L., S.C.M., C.M.M, K.M., and L.J.S.

## Acknowledgements

We thank the Institut Congolais pour la Conservation de la Nature (ICCN) for granting permission to conduct fieldwork on bonobos in the Salonga National Park buffer zone, the people of Lompole for facilitating research on bonobos in their forest, and Tanzania National Parks, the Tanzania Wildlife Research Institute, and the Tanzanian Commission for Science and Technology for granting us permission to conduct fieldwork on chimpanzees in Gombe National Park. We are extremely grateful to our many local collaborators at the LuiKotale Bonobo Project and the Jane Goodall Institute’s Gombe Stream Research Centre, particularly Lambert Booto, Mara Etike, Dr. Deus Mjungu, Madua Musa, Dismas Mwacha, and Sampson Pindu. We thank Joel Bray, Dr. Ian Gilby, and Dr. Kaitlin Wellens for contributing videos/photographs from which data for the current study were collected, and Zoe Andris, Sarah Fierman, Rosalie Mattiola, and Akhil Vedere for their assistance in screening and extracting still images from video footage. We also thank The George Washington University Primate Behavioral Ecology Lab and Primate Life History Lab for helpful feedback.

## Supplementary Information

**Supplementary Table 1.** Present/absent ranges

|  | Tooth | LuiKotale |  | Gombe |  |
| --- | --- | --- | --- | --- | --- |
|  |  | Oldest age<br>absent (days) | Youngest age<br>present (days) | Oldest age<br>absent (days) | Youngest age<br>present (days) |
| Maxillary | i1 | 85 | 88 | 120 | 261 |
|  | i2 | 85 | 123 | 120 | 261 |
|  | c | 477 | 227 | 120 | 571 |
|  | p3 | 85 | 123 | — | 526 |
|  | p4 | 85 | 227 | — | 526 |
| Mandibular | i1 | 85 | 88 | — | 120 |
|  | i2 | 117 | 123 | 123 | 175 |
|  | c | 474 | 227 | 261 | 371 |
|  | p3 | 117 | 123 | 120 | 261 |
|  | p4 | 123 | 216 | 120 | — |

“—” indicates that no score of present or absent was made for the given tooth.

**Supplementary Table 2.**
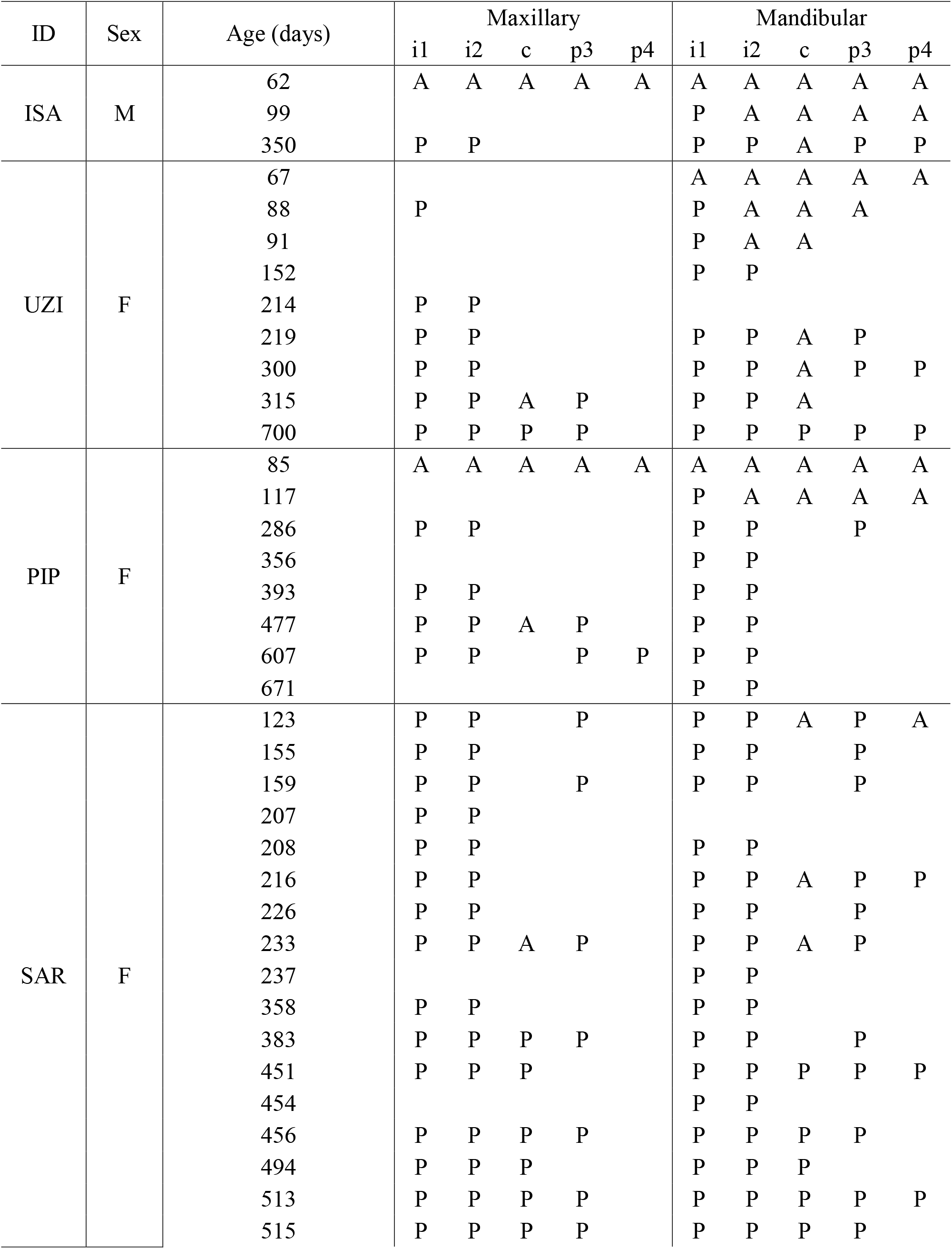
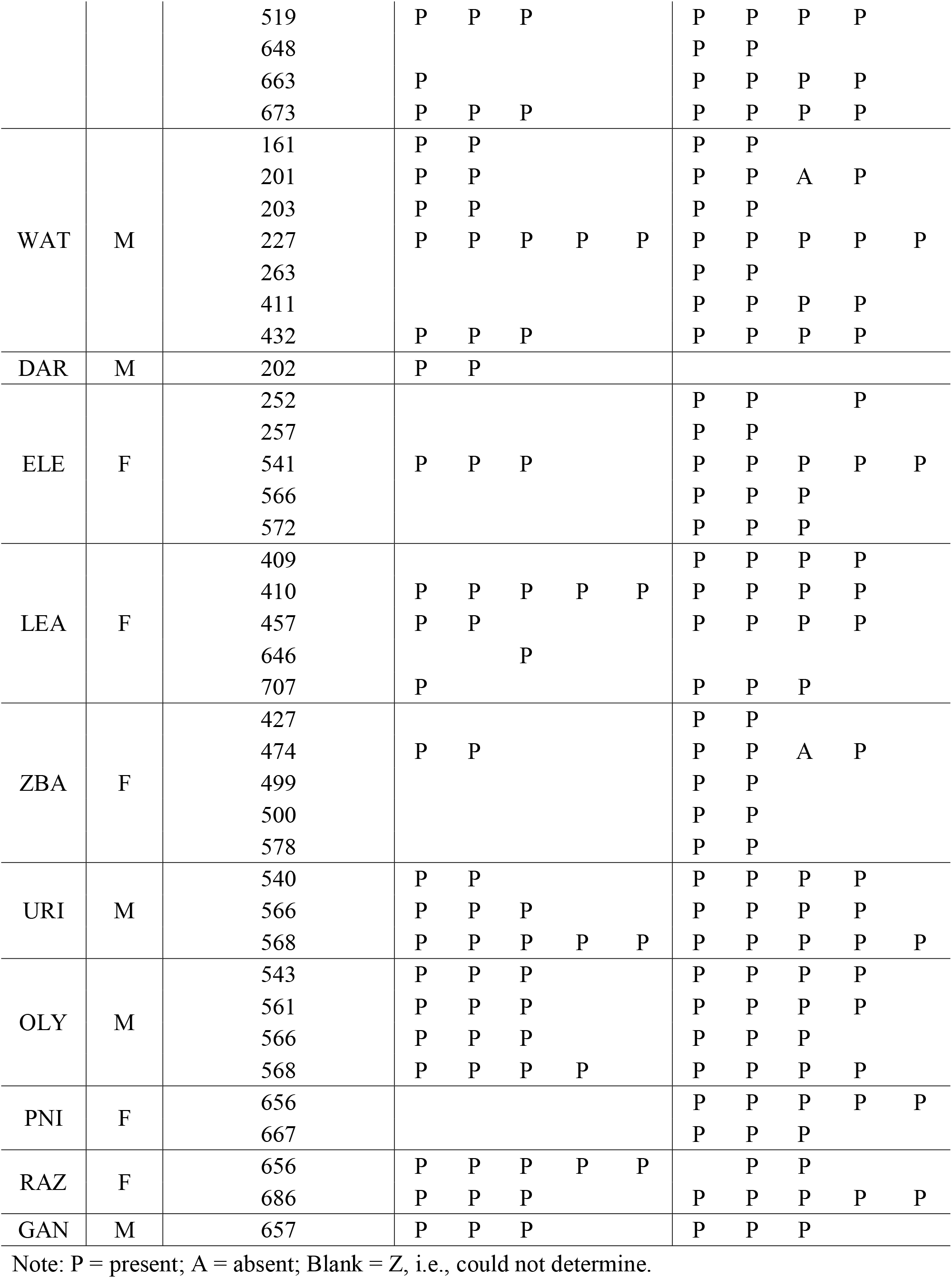
Aggregated raw scores for LuiKotale

**Supplementary Table 3.**
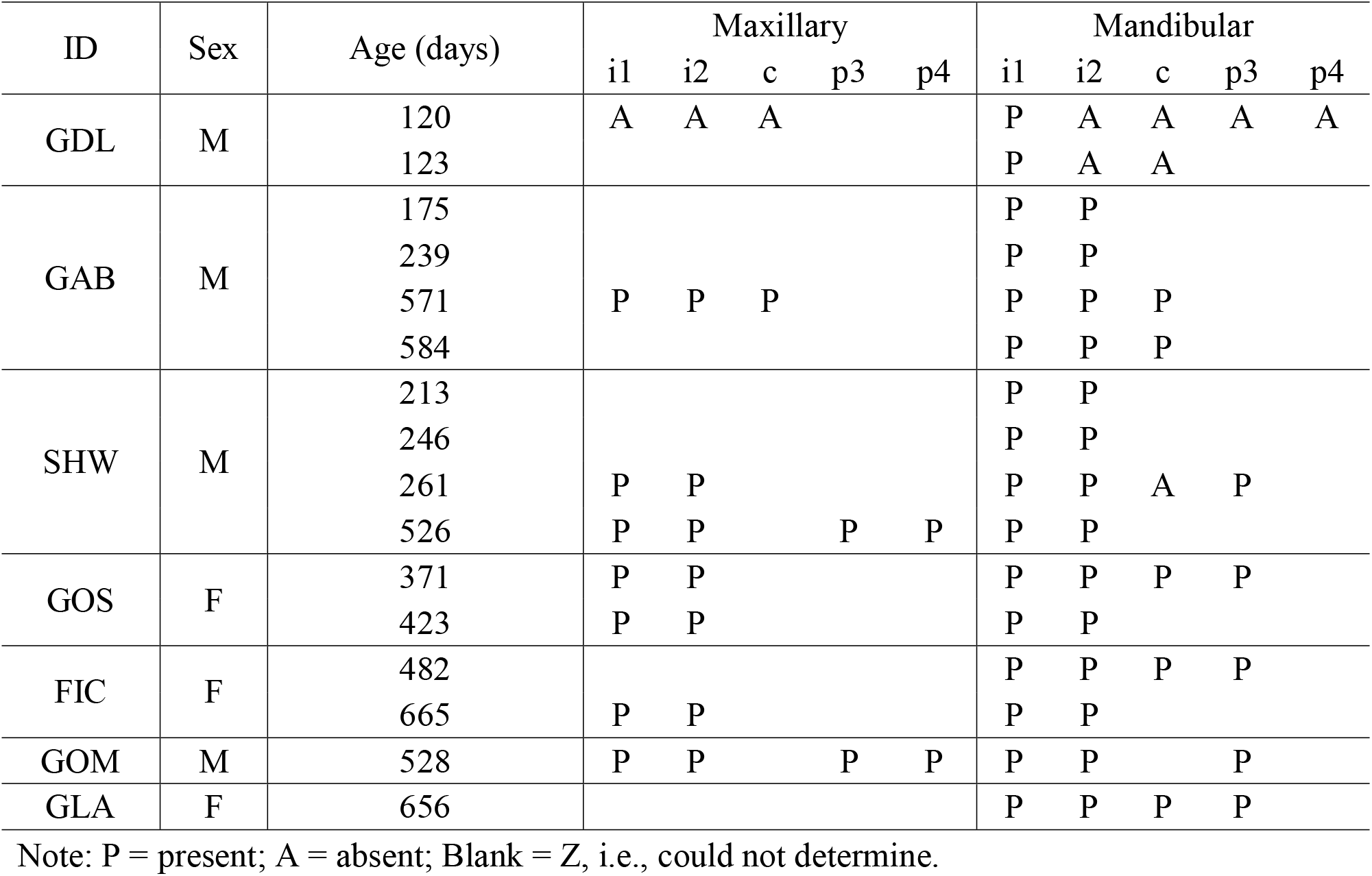
Aggregated raw scores for Gombe

